# Multi-site MRI analysis of morphometric differences in brain regions in the presence of hearing loss and tinnitus across the adult lifespan

**DOI:** 10.64898/2026.03.06.710136

**Authors:** Ivan Abraham, Shagun Ajmera, Wenhao Zhang, Amber M. Leaver, Bradley P. Sutton, Jonathan E. Peelle, Fatima T. Husain

## Abstract

The impact of age and hearing loss on the brain has garnered significant attention, as both factors have been implicated in the development of cognitive impairment or dementia. In this study, we investigated the impact of hearing loss and tinnitus on gray matter in the brain, while accounting for age. We used a comprehensive secondary analysis of structural MRI data obtained from multiple research sites (256 unique individuals) using voxel-based and surface-based morphology. After harmonization of this multi-site brain data, our research replicated the previously reported finding of age-related decline in total cortical volume, but there was no significant effect of either hearing loss or tinnitus on total cortical volume. When a region of interest analysis was conducted, the hippocampus emerged as the only brain region that showed a direct impact of hearing loss, after accounting for variance associated with age. This effect on hippocampal volume was evident in our sample from age 52 years onwards; when adjusted for hearing loss, the decline began at age 56 years. For the presence of tinnitus, ventral posterior cingulate gyrus showed main effects with respect to cortical volume and surface area while medial occipito-temporal gyrus and operculum of the inferior frontal gyrus showed significant main effects only with surface area. Post-hoc analysis revealed that posterior cingulate gyrus showed significantly higher volume and larger surface area in individuals with tinnitus compared to those without tinnitus. Similarly medial occipito-temporal gyrus surface area was increased whereas surface area of the inferior frontal opercular gyrus was reduced in those with tinnitus when compared to those without tinnitus. Notably, while past studies have reported that the presence of tinnitus appeared to moderate some of these effects in certain participant groups, our results suggest a more complex relationship between sensory degradation, chronic tinnitus, and brain structure in individuals across the adult lifespan.

**Highlights:**

- Hearing loss and tinnitus can exacerbate regional brain atrophy in the adult lifespan.
- High-frequency hearing loss affects auditory cortex gray matter volume to a larger degree in older age.
- Hearing loss may accelerate decline in hippocampal volume by about 4 years.
- Chronic subjective tinnitus is associated with a larger volume of cingulate cortex, increased surface area in cingulate cortex and the lingual gyrus, and decreased surface area of frontal operculum compared to controls.
- Tinnitus-related effects on regional brain atrophy are not modified by the degree of hearing deficits.

## Introduction

The human brain undergoes significant structural changes throughout the lifespan, with both cortical and subcortical regions exhibiting varying degrees of volume reduction as part of the normal aging process. These structural alterations have significant implications for sensory and cognitive functions in older adults. Research indicates that domain-specific patterns of regional brain atrophy contribute to individual differences in cognitive aging.

Magnetic Resonance Imaging (MRI) studies have consistently demonstrated that volumetric changes are not uniform across all brain regions. For instance, longitudinal analyses have revealed that critical structures such as the frontal and temporal lobes and the hippocampus experience pronounced atrophy with advancing age (Cardenas et al., 2011, Hedman et al., 2011, Raz et al., 2010). Specifically, the hippocampus, essential for memory formation, shows notable volume loss, correlating with age-related cognitive decline (e.g. Mungas et al., 2005). Conversely, certain regions like the cingulate gyrus and occipital cortex surrounding the calcarine sulcus appear more resistant to age-related gray matter density reductions (Raz et al., 2005). Further, volumetric reductions in the insula and superior parietal gyri have been associated with declines in specific cognitive domains, such as attention and visual-spatial abilities (Hedman et al., 2011).

Studies have shown that the rate of brain atrophy varies among individuals, even within the same age group, suggesting that factors beyond chronological aging, such as lifestyle and vascular health, play crucial roles in brain volume preservation (Fujita et al., 2023, Fletcher et al., 2016). Understanding these nuanced patterns of brain volume changes with age is vital for developing targeted interventions aimed at mitigating cognitive decline and promoting healthy brain aging. In this respect, understanding brain changes associated with age-related sensory decline is an important topic.

Sensory changes with age, particularly hearing loss, have been shown to have a significant impact on gray matter (Peelle et al., 2011, Eckert et al., 2019). Peelle et al. found that gray matter volume reduction in the primary auditory cortex (bilateral superior temporal gyri) was associated with poorer hearing thresholds, and reduced neural activations in the thalamus and brain stem were correlated with hearing decline. Eckert’s longitudinal study reported that higher high-frequency hearing thresholds were linked to lower gray matter volume in auditory cortical regions as well as greater expansion of lateral ventricles, which indicates a broader structural change in the brain beyond cortical thinning. Though similar patterns of results have been reported by larger-scale analysis – c.f. Kirschen and Leaver, 2024 or Zainul Abidin et al., 2025, these have lacked pure-tone audiometry, the gold standard assessment of hearing loss.

The present study investigates the impact of age, hearing loss and comorbid tinnitus using multi-site MRI data. Age and hearing loss are highly correlated and while the impact of each of these factors has been noted, their combined effect on brain morphology has not been documented as of yet nor has their effect in the presence of other hearing disorders (Lin, 2024). In a recent widely-cited study (Livingston et al., 2024), hearing loss was identified as the most relevant modifiable risk factor in middle-aged adults for developing dementia.

One important aspect of the current study is the inclusion of the disorder of tinnitus, which is often comorbid with age-related hearing loss. Tinnitus is the perception of noise or ringing in the ears without an external sound source (Biswas et al., 2022). It affects approximately 15-20% of people, with around 40-50 million individuals experiencing some form of tinnitus in the United States alone (Shargorodsky et al., 2010). This condition can result from exposure to loud noises, ear injuries, or circulatory system disorders and is often co-morbid with hearing loss. The economic burden of tinnitus is substantial, with estimated healthcare costs and lost productivity totaling around USD ∼30 billion annually in the U.S. (Trochidis et al., 2021). Surprisingly, given its prevalence, effective treatments vary, focusing on managing symptoms rather than curing the condition. Furthermore, the occurrence of tinnitus increases with age (Jarach et al., 2022) and the effect of tinnitus may not be additive but in some cases may moderate the effect of hearing loss on baseline age-related changes (Khan et al., 2021, Koops et al., 2020).

Despite this status quo, few large-scale studies of hearing loss and aging have taken tinnitus into account. The intersection of tinnitus and aging is clearly an important avenue to be studied but apart from a few studies (Husain and Khan, 2023, San-Martín et al., 2025), they have not been systematically investigated together. This work is the first attempt at addressing that gap using structural measures of brain regions from a large-scale multi-cite MRI dataset. Naturally, studies with small sample sizes, having low statistical power, may not lend themselves to simultaneous mapping of the impact of age, hearing loss and tinnitus.

Here, we have combined existing data from previous studies across five centers, leading to 265 unique participants. The data were acquired at multiple sites from distinct groups of individuals using different MRI scanners and acquisition parameters. Participants range in age from 18 to 81 years, with detailed hearing profiles on ∼72% and partial information on the remaining 28%. Using uniform data preprocessing pipeline for the raw MRI scans from all sites, and performing data harmonization with state-of-the-art techniques (Fortin et al., 2017), in a retrospective analysis, we investigated the individual contributions and the interplay between hearing loss, tinnitus, and age in brain morphology across the adult lifespan.

## Methods

### Participant data

Data collection was performed independently at each site: Washington University at St. Louis (WASHU), University of Illinois, Urbana-Champaign (UIUC), Wilford Hall Ambulatory Surgical Center (WHASC), Northwestern University (NU) and University of California - Los Angeles (UCLA). For each site, the data collection protocols were independently designed; however structural MRI data, pure-tone audiometry data, and status of chronic subjective tinnitus (tinnitus present for more than a year) were available for the large majority of individuals across the sites. Details of data used in this study are summarized below:

● **WASHU** (Rogers et al., 2020): Participants were among younger and older adults (n=60) and self-reported no history of neurological impairments or difficulties in hearing. Audiograms based on pure-tone audiometry were available for a subset of participants. Tinnitus status was not recorded for any participants. T1-weighted structural brain scan was acquired using an MPRAGE sequence (repetition time [TR] = 2.4 s, echo time [TE] = 2.2 ms, flip angle = 8°, 300 × 320 matrix, voxel size = 0.8 mm isotropic) on a 3T Siemens Prisma scanner.
● **UIUC** (Kim et al., 2025, Jain et al., 2024): Participants (n= 84) included one group having tinnitus (with a range of tinnitus severities) and a control group with no tinnitus. Both tinnitus and control groups had subgroups based on hearing loss profiling. In this study, we used only the tinnitus status information of individuals (i.e. binary variable indicating present/absent), and not the severity. Pure-tone audiometry was available for the low and high frequencies considered in this study. Based on Beck Inventory (Beck et al., 1988, Beck et al., 1996) none of the participants had severe depression or anxiety. T1-weighted images were acquired on a 3T Siemens Magnetom Prisma MRI scanner with the following parameters: TR = 2,300 ms, TE = 2.32 ms, flip angle = 8°, 192 slices, voxel size = 0.9 × 0.9 × 0.9 mm3.
● **WHASC** (Jain et al., 2024): Participants (n=74) were military-affiliated individuals, belonging to one of two groups, with tinnitus and without tinnitus. Both the groups included individuals with hearing loss and normal hearing. Pure-tone audiometry was available for the low and high frequencies considered in this study. Based on Beck Inventory (Beck et al., 1988, Beck et al., 1996) none of the participants had severe depression or anxiety. T1-weighted images were acquired on a 3T Siemens Prisma MRI scanner with the following parameters: orientation = sagittal, TR = 8.6 ms, TE = 4.00 ms, flip angle = 20°, voxel size = 1.2 x 1.1 x 7.0 mm³.
● **NU & UCLA :** These datasets included participants with and without tinnitus recruited at both NU (n=34) and UCLA (n=13) for previous studies (Leaver et al., 2022; Leaver 2025). Pure-tone audiometry was available for majority of these participants for the frequencies considered in this study. some volunteers had mild-to-moderate depression. T1-weighted images were acquired on 3T Prisma scanner with the following parameters: 0.8 mm isotropic, TR=2.5 s, TE=1.8, 3.6, 5.4, 7.2 ms combined, 1000 ms inversion time; 8° flip angle.

Age, sex and tinnitus status related statistics are summarized in Figure 1.

**Figure 1:**
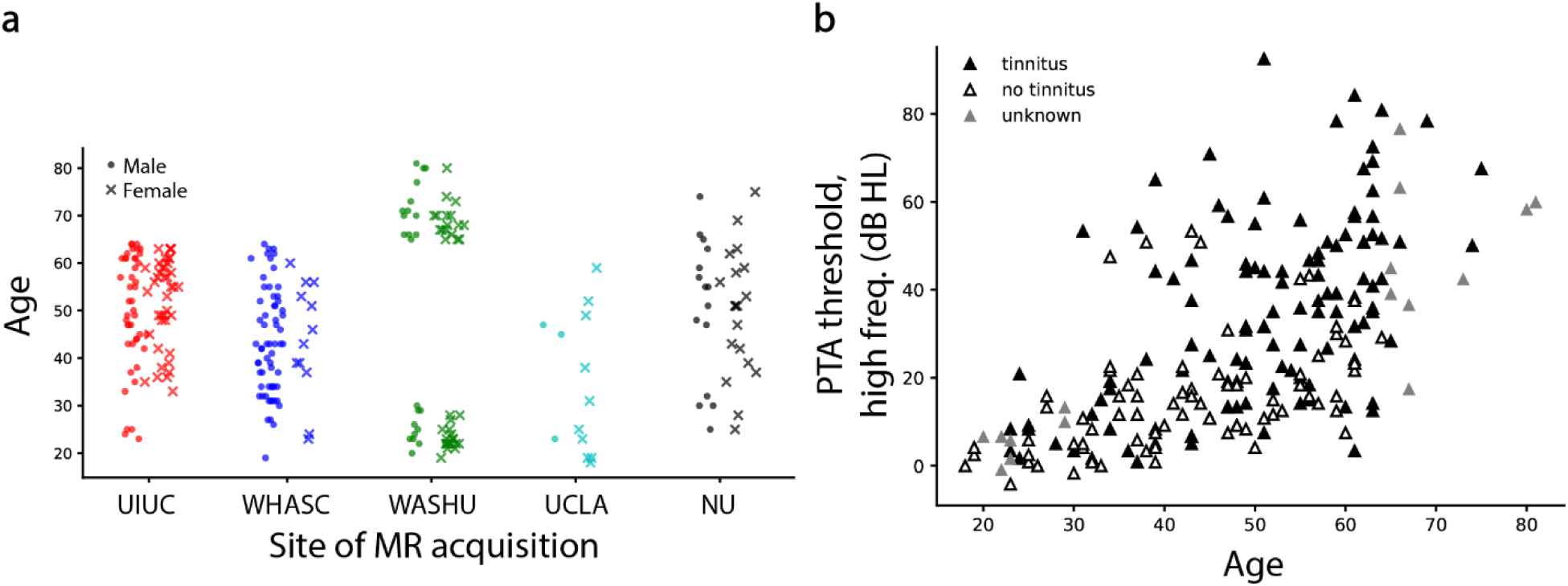
(a) Age distribution of individuals (y-axis) across sites where MRI data was acquired (x-axis); total n = 265. Individuals are shown separately by sex (M: dot markers; F: ‘x’ markers) **(b)** High-frequency pure-tone audiometry thresholds (y-axis) by individual age (x-axis), shown for n=212 individuals where PTA_high data was available. Participants’ tinnitus status are indicated by the markers described in the legend.

### Preprocessing Pipeline

The multi-site raw dataset consisted of T1- and T2-weighted brain images that were processed and analyzed uniformly across sites using standardized fMRI pre-processing and analysis routines available with the fMRIPrep software (Esteban et al., 2019). It must be noted that fMRIPrep-based pipeline is a standard preprocessing method in the field when analyzing relationships between individuals’ age and anatomical brain measures (Antonopoulos et al, 2023).

For each participant, across all sites, the T1-weighted (T1w) images were corrected for intensity non-uniformity (INU) with N4BiasFieldCorrection routine distributed with ANTs 2.3.3 and used as T1w-reference throughout the workflow (Avants and Gee, 2004). The T1w-reference was then skull-stripped using OASIS30ANTs as the target template. Brain tissue segmentation of cerebrospinal fluid (CSF), white-matter (WM), and gray matter (GM) was performed on the brain-extracted T1w image. Brain surfaces were reconstructed using recon-all routine in Freesurfer v.6.0.1 (Dale et al., 1999), and the brain mask estimated previously was refined with a custom variation of the method to reconcile ANTs-derived and FreeSurfer derived segmentations of the cortical gray-matter. Volume-based spatial normalization to two standard spaces (MNI152NLin2009cAsym, MNI152NLin6Asym) was performed through nonlinear registration with antsRegistration (ANTs 2.3.3), using brain extracted versions of both T1w reference and the T1w template. The following templates were selected for spatial normalization: ICBM 152 Nonlinear Asymmetrical template version 2009 [TemplateFlow ID: MNI152NLin2009cAsym] and FSL’s MNI ICBM 152 non-linear 6th Generation Asymmetric Average Brain Stereotaxic Registration Model [TemplateFlow ID: MNI152NLin6Asym].

### Morphological measures

The morphometric brain variables analyzed were regional cortical volume, regional surface area, and regional mean thickness. These measurements were available for 74 segments (sulci and gyri) each in left and right hemispheres, segmented as per Destrieux atlas (Destrieux et al., 2010); we summed between hemispheres the volume, area, and thickness measures for each region. Additionally, morphometric data for 27 other non-cortical brain components were also extracted from the aseg.stats file output by the preprocessing pipeline - these included the subcortical areas Amygdala, Hippocampus, Pallidum, Putamen, Caudate, Accumbens, Thalamus-Proper, Cerebellum-Cortex, Cerebellum-White-Matter, and sections of the corpus callosum and the ventricles. Other whole brain measures available for the participants were total cortical volume (TCV) and total intracranial volume (TIV). In total, 251 morphological variables were anaylzed.

To account for variability in individual brain size, we normalized the indices of regional cortical volume with total intracranial volume, and regional surface area with total surface area. We did not normalize the regional cortical thickness measures, as previous evidence with other cognitive disorders suggested doing so accorded no benefits (Westman et al., 2012).

### Data Harmonization

Because the participant data were acquired across several sites and scanners, we harmonized the dataset of anatomical variables for source bias (‘batch’ effects) using the neurocombat package (Fortin et al., 2017). We used the site location as the categorical ‘batch’ variable; age and sex were given as the biological variates to be preserved. Normalized measures of total cortical volume and total surface area, and average cortical thickness across regions, were compared with age across individuals, using Pearson’s correlation, both before and after data-harmonization. The effect of harmonization was measured with Cohen’s d effect size and was also analyzed using the Wilcoxon signed-rank test.

### Analysis - ANOVA With Anatomical Variables

After harmonization, we ran ANOVA for the families of variables of regional volume, regional surface area, and regional mean thickness. The volume and area measures were normalized for individual brain size (TIV and total surface area respectively); all variables were z-scored to facilitate post-hoc comparison of coefficients across regions. Pure-tone audiometry thresholds (PTA values) at 0.5, 1 and 2 kHz were averaged together to create a new variable PTA_low and PTA values at 4, 6 and 8 kHz were averaged together to form PTA_high. Tinnitus status was numerically encoded as 1, -1, and NaN for tinnitus present, absent and unknown status respectively. We used ordinary least squares (OLS) regression framework and Type-1 (sequential) ANOVA on each of the variables with the following models:

● 1 + age * PTA_low,
● 1 + age * PTA_high,
● 1 + age * tinnitus,
● 1 + age + PTA_high * tinnitus,
● 1 + age + PTA_low * tinnitus, and
● 1 + age * PTA_high + age * PTA_low + tinnitus

where * expands into summation of individual terms and interaction terms, e.g: age * PTA_high expands into age + PTA_high + age × PTA_high.

Our primary factors of interest in the ANOVA results were the final term in each of the models: the interaction effect between age and the other factors. In the absence of a significant interaction for any dependent variable, we tested the main effect for the factor in the final term (tinnitus status or PTA). p-values for all factors of interest were adjusted for multiple comparisons using the Benjamini-Hochberg method (Benjamini and Hochberg, 1995).

Following the ANOVA, the effects identified as significant were further analyzed as follows:

● For significant interaction effects: The intercept and all other predictors in the model, *excluding* the interaction term, were regressed out of the regional structural measure using their respective model coefficient estimates. The regional measure was then plotted against the interacting factors and visually analyzed to determine the underlying trends.
● For significant main effects: The intercept and all other predictors in the model, *excluding* the factor of interest, were first regressed out of the structural measure using their respective model coefficient estimates. Significance of relationship between the regional measure and the factor was then assessed using correlation values (for the PTA value) or a one-sided Welch’s *t*-test (for tinnitus status).

### Piecewise Regression Analysis

The significant main effect of hearing loss (PTA_high) on hippocampus volume was further analyzed by taking volume as the predicted variable and age as the predictor. Piecewise regression model was fit (for example: Ai et al., 2025) with number of breakpoints set to 1. The same procedure was repeated with the volume measure adjusted for hearing loss, i.e. with PTA_high regressed out using previously described OLS fit coefficients. Breakpoints in age, which represented change in volume trend, were compared for the two cases of hippocampal volume, i.e. with and without factoring the effect due to hearing loss.

## Results

### Behavioral and demographic measures

The collective sample of participants from across the five sites comprised 265 individuals in total, with age ranging from 18 to 81 years (median = 49 y.o; sex: 147 male, 117 female, 1 unknown; site-wise split shown in Figure 1a.) Pure tone audiometry for the lower frequencies of 0.5, 1, and 2 kHz, was available for 219 participants; the average ‘PTA_low’ threshold value (Supplementary Figure 1), had mean = 15.6 dB HL and standard deviation = 10.3 dB HL. ‘PTA_high’ threshold (averaged across 4, 6, and 8 kHz) was available for 212 participants, shown in Figure 1b, with mean = 27.0 dB HL and standard deviation = 20.9 dB HL. Overall, 80 subjects reported no tinnitus; 125 subjects reported having chronic subjective tinnitus presence.

### Whole-brain structural measures

The participant samples were aggregated from across different sites and MR scanners, thus, batch effects in data acquisition stemming from site differences could potentially bias brain anatomical measurements (e.g. Chen et al., 2014; Hawco et al., 2018). Hence, as a precursor to further analyses, we harmonized the data to remove site-related effects in the structural measures.

Overall cortical volume of individuals before data-harmonization was highly correlated with age (Good et al., 2001), yielding Pearson’s r = -0.654, p<0.001, 95% bootstrap CI [-0.720 -0.579]. After harmonizing the regional measurement dataset for site-differences, the correlation between individual cortical volume and age remained similar, namely Pearson’s r = -0.668, p<0.001, 95% bootstrap CI [-0.731 -0.594]. The effect of harmonization on whole-brain cortical volume was statistically insignificant (Cohen’s d = 0.003; paired two-sided t-test p-value = 0.875). See Figure 2.

**Figure 2:**
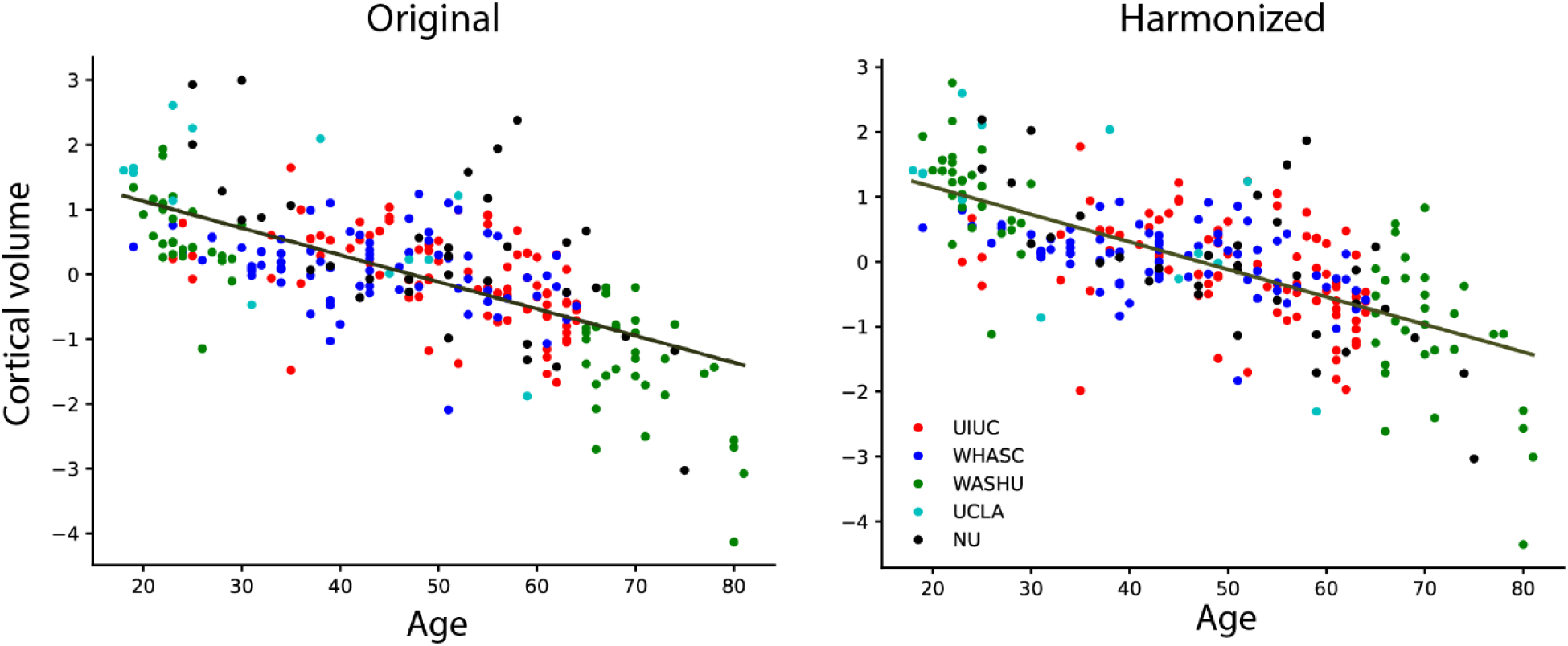
Y-axis: Cortical volume in the brain, normalized by individual total intracranial volume, and z-scored across participants. X-axis: individual age. Each dot represents a participant; color denotes the site of data acquisition. Lines depict linear fits to data. Left panel: original brain volume measures; Pearson’s r = -0.654, p=9.4e-34. Line fit: slope = -0.042, intercept = 1.960. Right panel: neuroCombat-harmonized brain volume measures; Pearson’s r = -0.668, p=1.3e-35. Line fit: slope = -0.042, intercept = 2.001.

Similarly, cumulative regional surface area and average cortical thickness also showed comparable correlations with age before and after harmonization. See Supplementary Figure 2 for details. The effects of harmonization were statistically insignificant for both total surface area (Cohen’s d = 0.001, and paired two-sided t-test p-value = 0.910) and average thickness (Cohen’s d =-0.004, and paired two-sided t-test p-value =0.859). Thus, the harmonization procedure removed site-specific effects without significantly altering the overall trends in cortical volume, thickness, and surface area measurements.

### Effect of hearing loss on region-wise anatomical features

After harmonizing across sites, we examined the structural brain indices for shared variance with individual hearing status. Pure-tone audiometry thresholds across high frequencies (PTA_high) showed a significant interaction with age across several brain measures (see Table 1), including gray matter volume of the transverse temporal sulcus (Figure 3a), gray matter volume of the amygdala, cerebellar white matter volume, global white matter volume, and lateral ventricle volume. Post-hoc analysis showed that, among individuals of comparable age, more severe hearing loss was related to further reduction in gray and white matter in the areas noted earlier and also increased enlargement of ventricles and axonal hypointensities (Figure 4).

**Figure 3:**
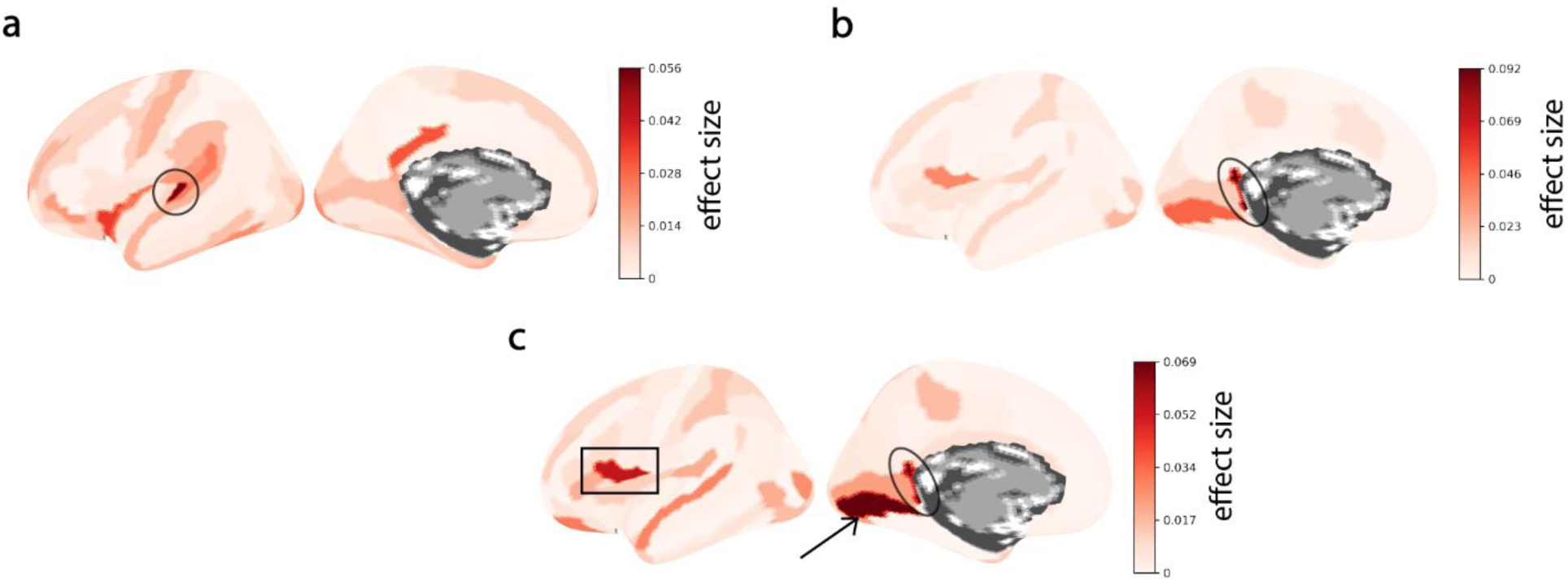
(a) Effect of interaction between age and hearing threshold on brain region volume. **(b)** Effect of tinnitus presence on brain region volume. **(c)** Effect of tinnitus presence on regional surface area. Colorbars show the effect size values (𝜂^2^) plotted on the lateral (left panels) and medial (right panels) brain surfaces. Only left hemisphere is shown. Regions highlighted are temporal transverse sulcus (circle), ventral posterior cingulate cortex (ellipse), inferior frontal operculum (rectangle), medial occipitotemporal cortex (arrow). See Tables 1 and 2 for other regions with statistically significant effects.

**Figure 4:**
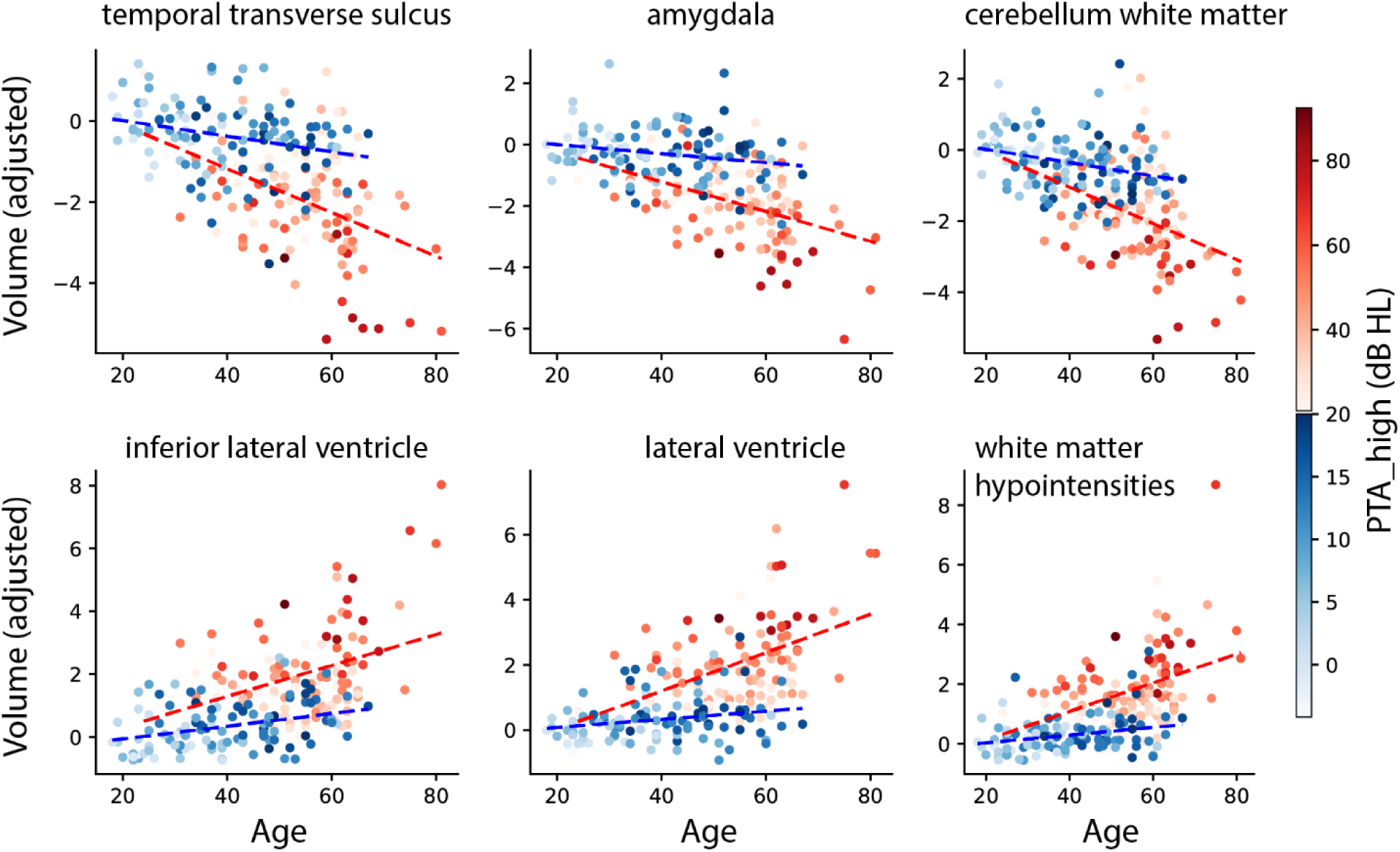
Visualization of interaction effect between age and hearing threshold on brain region volume. To facilitate visualization, PTA_high values are shown segregated by the mean value of 20.8 dB HL through red and blue colorscales and linear fits (dashed lines). Each dot represents a participant and the colorbar shows PTA_high values. Y axis: adjusted regional volume (z-scored), i.e., after factoring out the isolated (main) effects of age and hearing loss. The residual volume measures are plotted in the following order: auditory cortex (temporal transverse sulcus), amygdala, cerebellum white matter, inferior lateral ventricle, total lateral ventricle, and white matter hypointensities (see Table 1). High-frequency hearing deficits interact with age, inducing exacerbated decline in gray and white matter, and enlargement of ventricular cavities and white matter hypointensities.

**Table 1:** Volumetric differences in brain regions owing to hearing loss and tinnitus. Regions with Benjamini-Hochberg adjusted p-value<0.05 are shown. Region names: G_* indicates gyrus, S_* indicates sulcus. r: regional volume, a: age, p_hi_: PTA_high, t: tinnitus status, p_lo_: PTA_low

| Regional volume | Anova model | Factor | n | sum_sq | F-statistic | p-value (BH adjusted) | OLS fit coefficient |
| --- | --- | --- | --- | --- | --- | --- | --- |
| Lateral-Ventricle | r~1+a*p<sub>hi</sub> | a:p<sub>hi</sub> | 212 | 1.06E+01 | 15 | 1.43E-02 | 8.36E-04 |
| Inf-Lat-Vent |  |  |  | 1.02E+01 | 11.82 | 1.43E-02 | 8.20E-04 |
| S_temporal_transverse |  |  |  | 1.01E+01 | 12.44 | 1.43E-02 | -8.14E-04 |
| Amygdala |  |  |  | 9.11E+00 | 12.03 | 1.43E-02 | -7.74E-04 |
| Cerebellum-White-Matter |  |  |  | 8.54E+00 | 11.15 | 1.68E-02 | -7.50E-04 |
| WM-hypointensities |  |  |  | 7.98E+00 | 12.34 | 1.43E-02 | 7.24E-04 |
| Hippocampus |  | p<sub>hi</sub> | 212 | 1.09E+01 | 15.33 | 1.24E-02 | 1.68E-02 |
| G_cingul-Post-ventral | r~1+a*t | t | 205 | 1.06E+01 | 14.01 | 2.39E-02 | 2.91E-01 |
| G_cingul-Post-ventral | r~1+a*p<sub>hi</sub>+a*p<sub>lo</sub>+t | t | 195 | 1.28E+01 | 19.03 | 2.14E-03 | 2.94E-01 |
| none | r~1+a*p<sub>lo</sub> | a:p<sub>lo</sub>, p<sub>lo</sub> | 219 | - |  |  |  |
|  | r~1+a*t | a:t | 205 |  |  |  |  |
|  | r~1+a+p<sub>hi</sub>*t | p<sub>hi</sub>:t |  |  |  |  |  |
|  | r~1+a+p<sub>lo</sub>*t | p<sub>lo</sub>:t |  |  |  |  |  |

**Table 2:** Effect of hearing loss and tinnitus on surface area of brain regions. . Regions with Benjamini-Hochberg adjusted p-value<0.05 are shown. Region names: G_* indicates gyrus, S_* indicates sulcus. r: regional volume, a: age, p_hi_: PTA_high, t: tinnitus status, p_lo_: PTA_low

| Regional surface area | Anova model | Factor | n | sum_sq | F-statistic | p-value adjusted (bh) | coefficient |
| --- | --- | --- | --- | --- | --- | --- | --- |
| G_oc-temp_med-Lingual | $r \sim 1 + a * t$ | t | 205 | 1.18E+01 | 13.74 | 2.01E-02 | 5.32E-01 |
| G_cingul-Post-ventral |  |  |  | 9.94E+00 | 11.13 | 3.74E-02 | 2.64E-01 |
| G_oc-temp_med-Lingual | $r \sim 1 + a * p_{hi} + a * p_{lo} + t$ | t | 195 | 1.16E+01 | 13.87 | 1.38E-02 | 2.80E-01 |
| G_cingul-Post-ventral |  |  |  | 1.11E+01 | 13.13 | 1.38E-02 | 2.75E-01 |
| G_front_inf-Opercular |  |  |  | 9.64E+00 | 10.69 | 3.16E-02 | -2.56E-01 |

Next, the main effect of the PTA-high threshold was observed for hippocampal volume (after confirming absence of an interaction effect with age. See Table 1, model: 1 + age × PTA_high). After adjusting for age, hippocampal volume showed an overall decline with increasing PTA-high threshold, with a Pearson’s r of -0.319, p<0.001. Regional measures of surface area and mean thickness did not show any significant effects associated with PTA thresholds.

The hippocampus was the only brain region that showed variance in volume directly associated with hearing loss beyond any variance associated with age. We performed additional analysis to examine the relationship between hippocampal volume and age in our study sample after factoring out the effects due to hearing loss (PTA_high); this revealed a shift in the volume trend around the age of 56 years (95% CI: 49-63) which is consistent with findings from previous studies on age-related effects in the general population (Bowie et al., 2024; Nobis et al., 2019) – see Supplementary Figure 3. However, keeping the effect of hearing impairment intact in the data, a shift in the trend of hippocampal volume decline appeared to accelerate at 52 years of age (95% CI: 43-60) in our study sample.

### Effect of tinnitus on region-wise anatomical features

Next, we analyzed whether any variance in brain anatomical measures may be significantly attributed to tinnitus presence, beyond the variance that was explained by age and hearing loss. We estimated the effect of tinnitus by modelling for tinnitus status in isolation (with age), and in combination with the hearing thresholds.

There was no interaction effect of PTA thresholds with tinnitus status. Moreover, regions that showed tinnitus-related effects without factoring-in PTA thresholds were also found to be significant when the PTA covariates were included in the model; this was true for both cortical volume (Table 1) and surface area measures (Table 2). Ventral posterior cingulate gyrus showed significant tinnitus-related main effects for cortical volume as well as surface area (Figure 3b & Figure 3c). In addition, the surface area for medial occipitotemporal gyrus and operculum of the inferior frontal gyrus also showed significant main effects for tinnitus presence (Figure 3c).

We then performed follow-up analyses where we adjusted the regional measures for age and hearing thresholds then compared the distributions between individuals with and without tinnitus. Posterior cingulate gyrus showed significantly higher volume (p = 1.02e-06; Figure 5a) and larger surface area (p = 1.19e-05; Figure 5b) in individuals with tinnitus compared to those without tinnitus. Similarly medial occipito-temporal gyrus surface area was increased (p = 2.36e-05; Figure 5c), whereas surface area of the inferior frontal opercular gyrus was reduced (p = 1.54e-04; Figure 5d) in those with tinnitus when compared to those without tinnitus. Lastly, regional measures of mean thickness did not show any significant interaction or main effects associated with tinnitus status.

**Figure 5:**
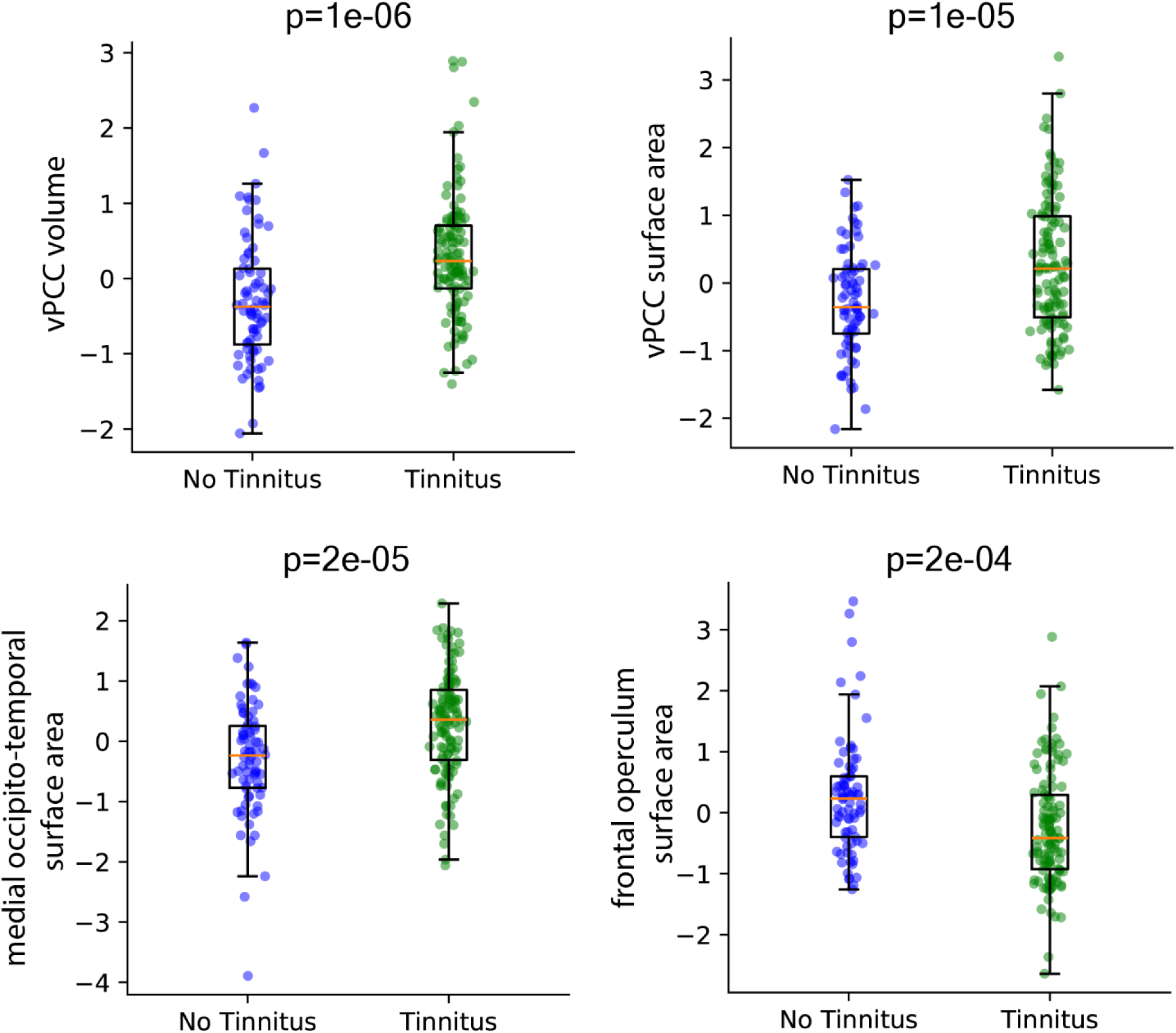
Effect of tinnitus on brain morphology. Comparison of regional measures (volume or surface area), after factoring our effects associated with age and PTA thresholds (low and high), between individuals with and without tinnitus (blue and green dots respectively). Comparison shown for **(a)** ventral PCC volume, **(b)** ventral PCC surface area, **(c)** medial occipito-temporal surface area, and **(d)** inferior frontal operculum surface area. One-sided t-test p-values are shown in the plots. Whiskers represent 1.5x inter-quartile range.

## Discussion

This study examined the effects of aging, hearing loss, and tinnitus on cortical and subcortical morphometry using harmonized structural MRI data from five independent sites. Our whole-brain regional analysis provides a comprehensive comparison through measurements of volume, surface area, and cortical thickness, as distinct patterns among these morphological measures may signify specific histological changes in the brain across adult age groups (Lemaitre et al., 2012). Across a diverse cohort of 265 individuals, we observed an age-related decline in global cortical volume, surface area, and thickness (Lemaitre et al., 2012). Furthermore, both high-frequency hearing loss and tinnitus were significantly associated with changes in regional cortical volume and surface area respectively, and we found evidence for an interaction between age and high-frequency hearing loss. Notably, while past studies have reported that the presence of tinnitus appeared to moderate these effects in some participant groups (Koops et al., 2020) – albeit in a much smaller sample sized study – our results suggest a more complex relationship between sensory degradation, tinnitus, and brain structure in aging individuals.

### Effect of hearing loss

Recall that indices of cortical and subcortical gray and white matter, as well as ventricular spaces, were examined for alterations associated with individuals’ hearing thresholds. A reduction in hippocampal gray matter volume was significantly associated with the degree of high-frequency hearing loss (HFHL), independent of age-related differences, in accordance with the results from a previous large-scale VBM study (Shim et al., 2023).

High-frequency hearing loss also appeared to differentially affect cortical volume decline in the auditory cortex with age and was associated with increased lateral ventricle expansion. Age-related hearing loss likely affects brain structure through two separate mechanisms (Eckert et al., 2019). Eckert’s study previously noted two statistically independent changes in brain morphology associated with age-related hearing loss. Their cross-sectional analysis consistently showed that elevated HFHL was associated with lower gray matter volume in the auditory cortex though volume changes did not track with changes in hearing over time. At the same time, participants with greater HFHL at the study’s onset exhibited a significantly faster rate of lateral ventricle volume expansion over time and the authors concluded that age-related hearing loss likely affects brain structure through two separate pathways.

Similar interactions between age and hearing function were also reported for inferior lateral ventricle volume, auditory cortex thickness, and other regions in a previous study (Kirschen and Leaver, 2024); however, this study used a much coarser measure of hearing function (words-in-noise task) compared with pure-tone audiometry used in the current study and in Eckert et al.. Additionally, hearing impairment in older adults was associated with reduced global white matter volume across the brain (Rigters et al., 2017). This association was found to be consistent across different hearing frequencies (low and high) and was found to be independent of common risk factors like age, sex, cognitive function, and cardiovascular risk factors.

### Effect of tinnitus

Individuals with chronic tinnitus typically exhibit altered resting-state functional connectivity in regions implicated in self-awareness, such as the precuneus and posterior cingulate cortex (PCC) (Schmidt et al., 2017; Schmidt et al., 2013). Our findings align with these previous studies by highlighting tinnitus-related abnormalities in ventral PCC and retrosplenial cortex morphometry, which could be closely linked to the altered functional involvement of these regions (Schmidt et al., 2017). Makani et al also report in their meta-analysis that individuals with chronic tinnitus show significantly increased gray matter levels in the bilateral lingual gyrus that was dependent on matching for hearing-loss (Makani et al., 2022). They interpret this as follows: the co-occurrence of tinnitus with hearing loss is associated with preserved gray matter in the lingual gyrus (and precuneus) compared to individuals with hearing loss alone, suggesting that hearing loss is the major driving factor of gray matter changes, and tinnitus may counteract some of the age- or hearing-loss-related decline in this region.

The limbic system has been previously implicated in multiple studies involving tinnitus – both in terms of functional connectivity (e.g. Zimmerman et al., 2018) as well as in terms of morphology (e.g. Profant et al., 2020). Relevant to the present study, Profant et al. reported in their study (n=73) that tinnitus was associated with an increase in the volume of the amygdala but presbycusis had no significant association nor did tinnitus severity (Tinnitus Handicap Inventory score). However, in the present study, we found that a decreasing trend in amygdalar volume associated with aging but not due to tinnitus presence, possibly due to the ameliorative effect of tinnitus.

We observed reduced surface area in the inferior frontal opercular gyrus in those with tinnitus compared to controls. This area is typically engaged with speech production and part of the canonical language network, and with cognitive control (Hickok, 2022). In a subset of the participants from the UIUC cohort, those with normal hearing thresholds and tinnitus, we have previously related the decline in gray matter volume in the inferior frontal gyrus to performance on the Speech-in-noise (SiN) task (Tai et al., 2023). We observed decreased volume in the IFG relative to age- and gender-matched controls, but this was not correlated with the SiN performance. Instead, the SiN performance negatively correlated with GM volume in the left cerebellum (Crus I/II) and the left superior temporal gyrus. An earlier study by Wong et al., contrasting older adults with mild hearing loss and younger adults with normal hearing thresholds, had noted that in general, there was reduced GM volume across several different regions of interest, which the investigators attribute to age-related atrophy (Wong et al., 2010). When regional brain volumes were correlated with SiN performance, only regions in the prefrontal cortex, left pars triangularis and the cortical thickness of the left superior frontal gyrus, were observed to be significant predictors of the SiN performance of the older participants. Understanding speech in noisy listening situations increases cognitive load, especially in the context of hearing loss (Peelle, 2018) or perceiving continuous tinnitus sound(s) (Husain and Khan, 2023). The pattern of atrophy noted above may be linked to such increased attentional and cognitive load.

Unsurprisingly, tinnitus (the perception of internally generated sound) and hearing loss (auditory deprivation) appear to exert distinct effects on neural tissue. Notably, tinnitus-related alterations in the aforementioned regions persisted even after controlling for high-frequency pure-tone thresholds. These findings underscore the necessity for neuroimaging studies to account for hearing loss when investigating tinnitus, and vice versa, given the high comorbidity of these conditions in the general population. Failure to adequately dissociate their contributions may lead to confounded neural markers in tinnitus research.

### Caveats and Future Directions

While the sample size of the combined dataset is an order of magnitude greater than typically found in VBM/SBM studies, an important consideration in this multi-site study is the potential variability introduced by differences in MRI scanners, acquisition protocols, and site-specific participant characteristics. To mitigate these sources of heterogeneity, all raw MRI data were processed using a standardized pre-processing pipeline that included denoising, correction for intensity inhomogeneity, skull stripping, etc. They were further harmonized using techniques introduced by Fortin. et al. This harmonized approach ensured consistency in cortical volume estimates across diverse imaging conditions. However, it is important to acknowledge that deeper scanner-specific biases or domain shifts—such as those arising from hardware differences, coil sensitivity profiles, or manufacturer specific variations—may still persist. These types of intrinsic and latent variability often require more advanced approaches, such as machine learning (ML) based domain adaptation (Guan et al., 2021) or out-of-distribution learning techniques (Mårtensson et al., 2020), which were beyond the scope of the current work. Future studies incorporating such strategies may offer improved sensitivity in disentangling biological effects from technical artifacts in multi-site neuro-imaging data.

While the study observes important cross-sectional trends in the brain across adult lifespan, it does not allow inferences about the progression in regional morphology within aging individuals owing to tinnitus and hearing degradation. Longitudinal brain data acquired at multiple time points along aging are needed to better attribute any changes in regional morphometry to the presence of hearing impairments (Fitzhugh & Pa, 2022, Slade et al., 2022).

## Conclusion

This study provides robust evidence for the independent and combined effects of age, hearing loss, and tinnitus on cortical gray matter volume across adulthood. By leveraging a harmonized, multi-site MRI dataset and uniform preprocessing methods, we demonstrated and confirmed the declining trends in cortical volume, surface area, and cortical thickness with age in our sample. We found a profound effect of hearing loss but an equivocal effect of tinnitus. This supports emerging evidence that tinnitus may interact with hearing loss in complex ways, potentially moderating the neuroanatomical impact of auditory decline. However, further investigation with larger, tinnitus-focused cohorts are needed to clarify this relationship.

The findings reported here also validate the utility of large-scale, combined neuroimaging datasets in characterizing structural brain changes across heterogeneous patient populations. Using toolboxes such as NeuroCombat allowed us to account for residual variability introduced by scanner and any site-specific differences. Taken together, our results emphasize that auditory health—particularly hearing loss—has a measurable impact on brain structure above and beyond normal aging. These findings have important implications for public health, as they highlight the potential for hearing-related interventions to mitigate cortical atrophy and its associated cognitive risks in aging populations. Future studies should aim to integrate additional clinical, cognitive, and lifestyle factors to better contextualize these structural changes and support the development of targeted, sex-sensitive therapeutic strategies.

## Supplementary

**Supplementary Figure 1:**
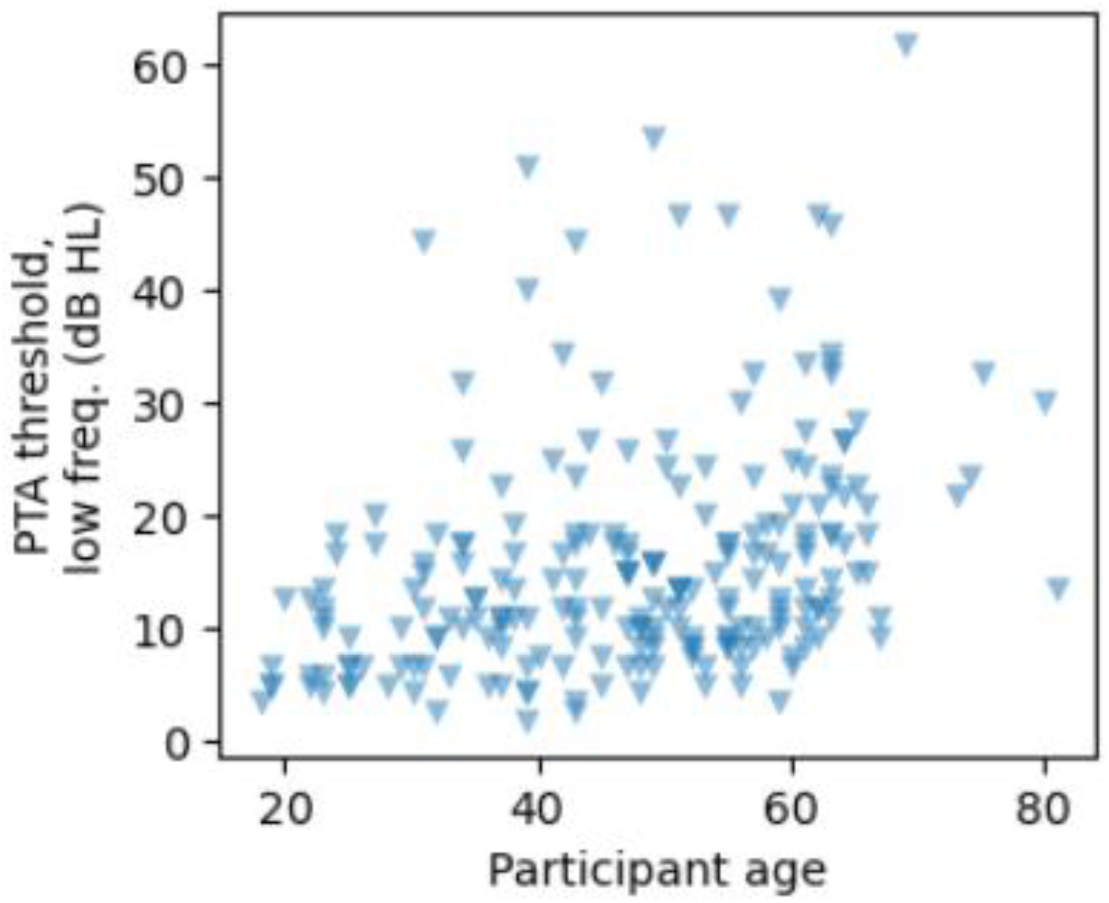
Low-frequency pure-tone audiometry thresholds (y-axis) by individual age (x-axis) shown for n=219 individuals where PTA_low data was available.

**Supplementary Figure 2:**
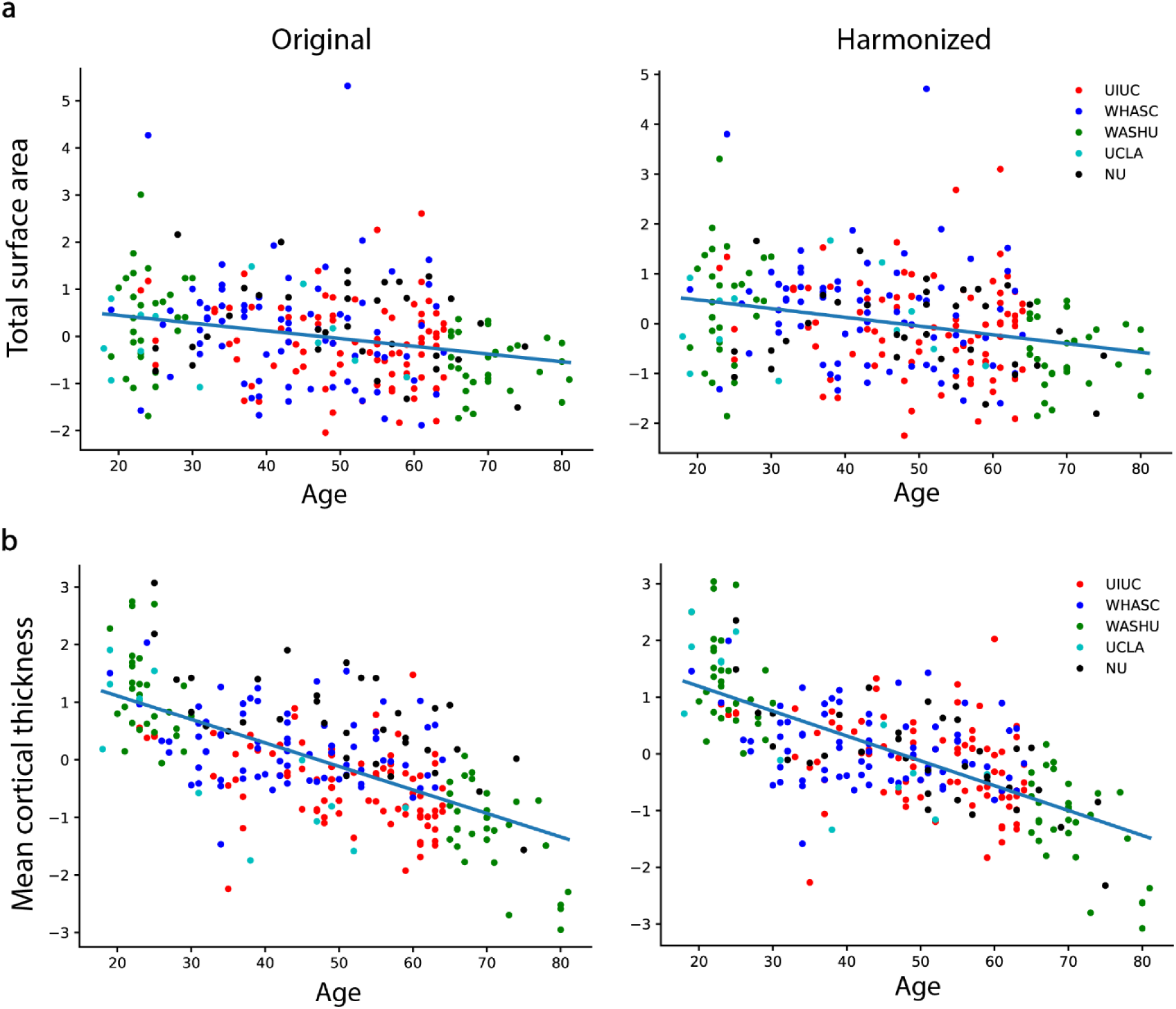
(a) Y-axis: *Brain surface area* z-scored across participants. X-axis : individual age. Lines depict linear fits to data. Left panel: original surface area measures. Pearson’s r = -0.258, p= 2.0e-05. Line fit: slope = -0.016, intercept = 0.774. Right panel: neuroCombat-harmonized surface area measures. Pearson’s r = -0.276, p= 4.9e-06. Line fit: slope = -0.018, intercept = 0.828. **(b)** Same as (a) but for average cortical thickness measure. Left panel: Pearson’s r = -0.643, p= 2.6e-32. Line fit: slope = -0.041, intercept =1.926. Right panel: Pearson’s r = -0.690, p= 7.9e-39. Line fit: slope = -0.044, intercept = 2.068. Each dot represents a participant; colors denote the site of data acquisition.

**Supplementary Figure 3:**
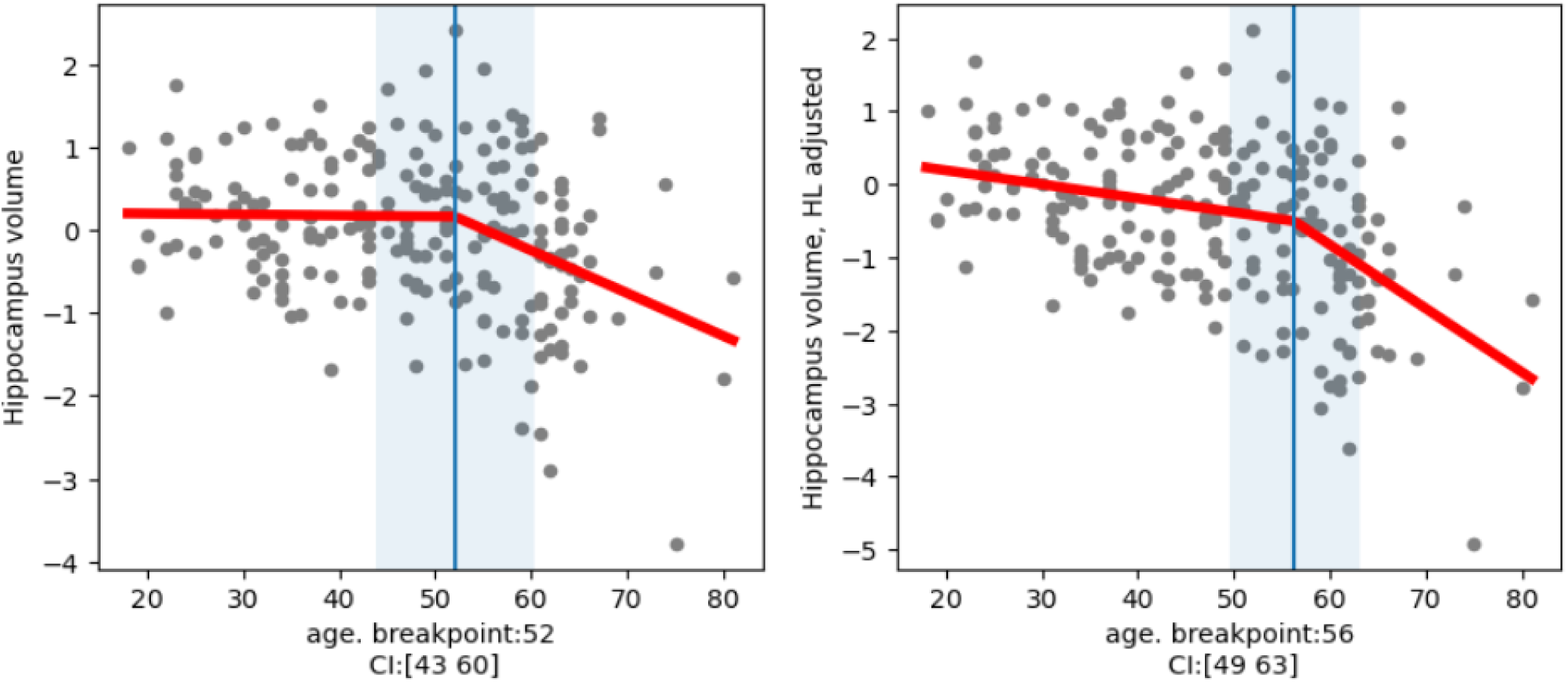
Left panel: Volume of bilateral hippocampus across all individuals in study sample (where PTA_high data was available; n=212). Right panel: same as left, but after factoring out effects of high-frequency hearing thresholds from the volume measure. Red lines represent piecewise linear fits to data. Vertical blue line: breakpoint in trend. Shaded blue region: 95% confidence interval for the estimated breakpoint. Left panel: slope1 = -0.001 [-0.019 0.017]; slope2 = -0.051 [-0.081 -0.022]. Right panel: slope1 = -0.019 [-0.035 -0.003]; slope2 = -0.087 [-0.132 -0.042].

